# Evaluating the Effectiveness of Parameter-Efficient Fine-Tuning in Genomic Classification Tasks

**DOI:** 10.1101/2025.08.21.671544

**Authors:** Daniel Berman, Daniel Jimenez, Stanley Ta, Brian Merritt, Jeremy Ratcliff, Vijay Narayan

**Author notes:** Co-senior authors (Jeremy Ratcliff, Vijay Narayan). Google DeepMind, 14–18 Handyside Street, London, N1C 4DN, United Kingdom (Jeremy Ratcliff).

## Abstract

Foundation models are increasingly being leveraged for biological tasks. To address the high memory requirements of fine-tuning large pre-trained language models, parameter efficient fine-tuning (PEFT) methods are also being increasingly utilized. Previous studies have shown minimal, if any, loss in performance when using PEFT on binary classification tasks. However, the impact of using PEFT on tasks with large classification spaces has not been systemically evaluated. In this work, we apply PEFT to the problem of taxonomic classification using pre-trained genomic language models as the classification backbone. We explore various training strategies—including PEFT, full fine-tuning, and partial fine-tuning—for classifying sequences at the superkingdom, phylum, and genus levels. We find that PEFT-trained models significantly underperform compared to those trained via full fine-tuning or partial fine-tuning. Additionally, we demonstrate increased performance of pretrained models over those randomly initialized.

## INTRODUCTION

Adapting techniques from natural language to biological sequence data may yield impactful performance across a variety of tasks.^1,2^ Building on advancements in large language models (LLMs), foundation genomic language models (gLMs) and protein language models (pLMs) are pre-trained on large corpuses of unlabeled data. These models can be applied to specific tasks through fine-tuning of pretrained weights; for example, fine-tuning has been repeatedly demonstrated to enable a wide-range of classification tasks on biological data.^3,4^

However, fine-tuning introduces significant memory requirements, both in terms of random-access memory (RAM) and storage, which is computationally expensive and limits adoption of these tools by compute limited laboratories. To mitigate storage problems associated with multiple fine-tuned models, parameter-efficient fine-tuning (PEFT) methods were devised, where only a fraction of the weights need to be trained and saved as a new model while maintaining the core model as the base.^5,6^ Therefore, N tasks no longer require N+1 models to be saved, but rather one full model and several sets of parameters that amount to fewer than 1% of the full model weights.

These PEFT methods, such as low rank adaptation (LoRA)^8^ and infused adapter by inhibiting and amplifying inner activation ((IA)^3^),^9^ have been adapted to gLMs and pLMs.^10-14^ However, these approaches have typically been applied to binary classification tasks, such as promoter/enhancer detection and signal peptide prediction. To evaluate the appropriateness of PEFT methods over a larger classification space, we systematically evaluated the performance of multiple training approaches in a hierarchal classification task with a progressively expanding classification space for fixed inputs. Here, we observe a significant reduction in performance when using PEFT compared to full fine-tuning as the classification space expands. In addition, we fail to observe improvements in runtime efficiency or memory usage.

## EXPERIMENT

In our experiments, we fine-tuned pre-trained gLMs for taxonomic classification using short nucleotide sequence fragments of fixed length (1500bp) at three taxonomic ranks. Four training methods were applied to five models across three architectures: Nucleotide Transformer (NT) 50M, 100M, and 250M,^3^ DNABERT-S,^15^ and DNABERT-2.^16^ Models were fine-tuned to predict superkingdom, phylum, and genus labels on a taxonomy dataset containing 4, 57, and 1,884 classes, respectively. Models were trained using the following strategies:

1. fine-tuned
2. fine-tuned after initialization to random weights.
3. LoRA (multiple parameters)
4. (IA)^3^
5. fine-tuned with 75% of model weights frozen

Models were trained for five epochs unless otherwise specified. We used a heuristic linear scaling to adjust learning rate with training batch size for each of the gLMs considered. We fine-tuned DNABERT-2 and DNABERT-S with initial learning rates of 2e-5, NT 100M and NT 250M with initial learning rate of 1e-4, and NT 50M with an initial learning rate of 2e-4. In addition, a cosine learning rate scheduler was used to adjust the learning rate during training. Models were trained using FP32 precision and the AdamW optimizer and initialized with the default pretrained weights available from HuggingFace, with the exception of the randomly initialized weights. PEFT methods were implemented using HuggingFace’s PEFT library (https://huggingface.co/PEFT). A complete list of experimental results is contained in the supplementary dataset. See extended methods for further details on model training, hyperparameter optimization for PEFT methods, and training and test sets.

## RESULTS AND DISCUSSION

### Classification Performance

We found that fine-tuned models consistently achieve >99% balanced accuracy when predicting superkingdom for the test set under consideration. In addition, balanced accuracy decreased as the classification space expanded for all models and training strategies considered, as expected. For each fine-tuned model, the balanced accuracy decreased by a median of 1.6% when comparing superkingdom to phylum. Balanced accuracy decreased a further median of 25.8% when comparing phylum to genus.

A recent study suggested that the weights assigned when pre-training gLMs such as NT^3^ and DNABERT-2^16^ do not translate to increased classification performance on a number of binary and 3-class classification tasks focused on epigenetics, promotors, and enhancers.^7^ To assess this claim, we fine-tuned a set of models whose initial weights were randomly initialized, obliterating any learning from model pre-training. In contrast to the observations described in Vishniakov et al. (2024), randomly initialized models underperformed fine-tuned models for every model and every rank at a magnitude as high as 22% on balanced accuracy (genus) for NT 50M (Figure 1a). The loss in performance was largest for the NT model architectures. This result is highly suggestive that model pre-training embeds some degree of knowledge as to the differences between taxonomic ranks and that performance gains from this practice are not marginal.

**Figure 1:**
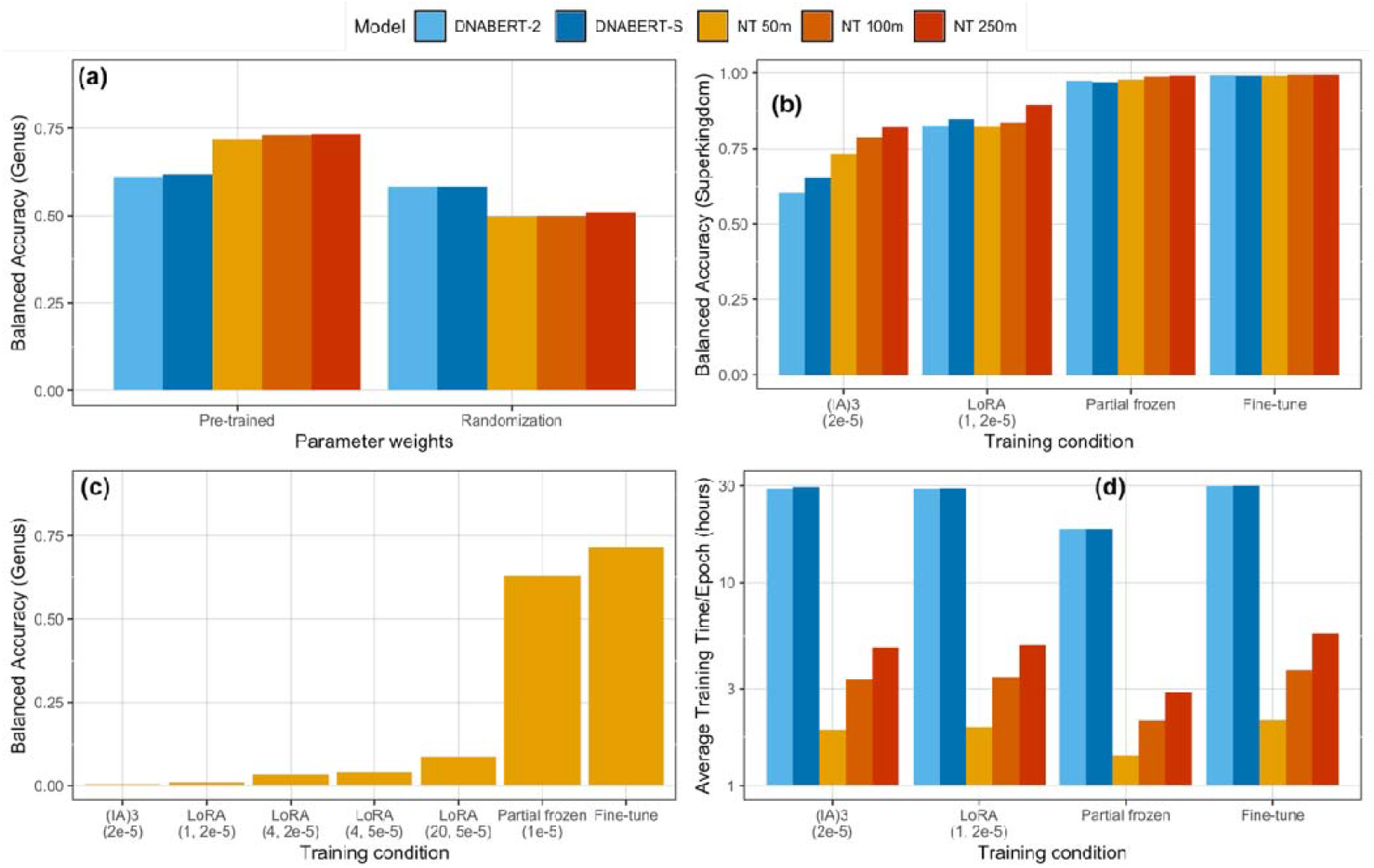
Model training results. (**a**) Balanced accuracy for phylum predictions of fine-tuned pre-trained models and fine-tuned models initialized with random weights. (**b**) Balanced accuracy for superkingdom prediction for models trained with PEFT, frozen layers, or fine-tuned. Parentheses record learning rate for (IA)^3^ and r and learning rate for LoRA. (**c**) Balanced accuracy for genus prediction of the NT-50M model with a variety of training approaches and variation in LoRA parameters. (**d**) Training efficiency for different training approaches, measured by average training time per epoch.

Models trained using PEFT methods substantially underperformed fine-tuned models across all ranks with the performance gap increasing as the classification space expanded (Figures 1b).

Models trained with (IA)^3^, for instance, had mean balanced accuracy scores 26% lower for superkingdom and 83% lower for phylum. A more in-depth investigation of the NT-50M model demonstrated that LoRA scores scaled as the parameter r increased but still grossly underperformed fine-tuning (Figure 1c) Models trained with 75% of layers frozen performed most closely to fine-tuned models, but were still less accurate for the larger classification spaces; decreases for superkingdom predictions ranged from 0.6% to 2.1% while those for genus ranged from 5.6% to 12.6%.

### Training efficiency

In addition to evaluating the impact of PEFT methods on the classification performance of various fine-tuned gLMs, we also examined their effect on training efficiency (Figure 1d). To do this, we explored two key factors: training time per epoch and batch size. While batch size and training time are typically inversely proportional, reducing trainable parameters could potentially free up memory, allowing for increased batch size and potentially accelerating training.

To find the maximum batch size, we configured a batch size of 4 and trained one epoch before doubling the batch size and repeating, until reaching the GPU memory limit. Training was conducted on a single NVIDIA H100 with 80GB of GPU memory for the NT models, and on 4 NVIDIA H100s for the DNABERT models.

It is important to note that while model performance is strongly influenced by the learning rate, training time per epoch does not directly depend on it. For this analysis, we scaled the learning rate linearly with the increased batch size with the aim of determining the maximum batch size enabled by PEFT methods regardless of convergence or performance.

Ultimately, we found that despite reducing the number of trainable parameters, the batch size limit remained unchanged across all tested configurations. As a result, we also found that neither LoRA nor (IA)^3^ provided any substantial benefit in terms of training time (Figure 1d). (IA)^3^ provided mild decreases in training time, ranging from 1.3% to 14.7%. LoRA, on the other hand, showed decreases of 2.8% to 12.2% when r=1 but increases of 6.1% to 22.4% when r ≥ 4. In contrast, training with partially frozen weights demonstrated consistently large decreases in training time from 32.4% to 48.7%.

## CONCLUSION

As machine learning based methods continue to be integrated into the analysis of biological data, intelligent selection of appropriate training methods is paramount. PEFT techniques such as LoRA and (IA)^3^ provide insufficient performance when applied to tasks with large classification spaces, such as taxonomic classification. These limitations are reflected in both model classification performance and overall training time. In addition, the results of this study contradict the conclusions of Vishniakov et al. (2024) and strongly suggest model pre-training can improve performance on predictive tasks, at least for taxonomic classification. Future work should focus on the limits of PEFT in other domains with large classification spaces, especially where a hierarchical nature of the labels exists, such as text classification or protein family classification.

## Supporting information

PEFT Supplementary Complete Results

## Code Availability

Model training and inference code used in this study is available at https://github.com/jhuapl-bio/microbert.

## Acknowledgements

This work was supported by funding from the U.S. Centers for Disease Control and Prevention (CDC) through the Office of Readiness and Response under Contract No. 75D30124C20202. We would like to thank Dr. Molly Gallagher for valuable contributions and for helping us navigate programmatic hurdles during the completion of this work.

## EXTENDED METHODS

### Taxonomic Data

The DNA sequence data used to fine-tune the gLMs in this analysis is sourced from the open-access dataset used for pre-training in the publication *“BERTax: Taxonomic Classification of DNA Sequences with Deep Neural Networks”*. The dataset is available as supplemental material: https://osf.io/dwkte. The available data consists of four separate FASTA files, each corresponding to one of the four superkingdoms of interest: Archaea_db.fa, Viruses_db.fa, Eukaryota_db.fa, and Bacteria_db.fa. All sequences are of fixed length—1500 base pairs. We preprocess these FASTA files by standardizing all sequences to uppercase. All the pre-trained gLMs considered have maximum token lengths that comfortably exceed the lengths of the input sequences. For example, the NT models use 6-mer tokenization with a maximum token length of 1000, which is significantly larger than the expected input size of 250 tokens for the sequences in our dataset.

To assign taxonomic labels, we use the ncbi_taxonomy module from ETE to retrieve Phylum and Genus names from provided taxonomy ID values.^17^ This module enables querying of a local copy of the NCBI Taxonomy database, which we built using the December 2024 version of the publicly available taxdump archive: https://ftp.ncbi.nlm.nih.gov/pub/taxonomy/taxdump_archive/. Sequences lacking either a phylum or genus label are excluded from our analysis.

The full processed dataset includes 5,181,880 sequences: 2,601,890 from Eukaryota, 1,828,018 from Bacteria, 524,276 from Archaea, and 227,696 from Viruses. These are annotated with a total of 1,573 unique taxonomy IDs, spanning 4 superkingdoms, 55 phyla, and 1,878 genera. We reserve 2% of the full dataset stratified by Genus labels (103,638 sequences) as a holdout test set. This test set is used to evaluate all experimental runs. We also reserve 10% of the full dataset as a validation set to monitor model performance by evaluating validation loss.

### Model Architectures

We developed custom hierarchical classification models using pre-trained gLMs as the backbone. After initializing the base model, we add a set of three classification heads, one for each taxonomic rank under consideration. Each head is implemented as a linear layer that maps the transformer’s hidden size to the number of class labels for that rank, and each of the classification heads is independent of each other. The implementation of a fully hierarchical architecture, where outputs from one rank inform the next, is left for future exploration.

During a forward pass, input token ids and attention masks are processed by the base model to produce token-level hidden states. We extract the last hidden state and apply mean pooling across the tokenized sequence to obtain a fixed-size embedding for each input (512 dimensional for NT-50M and NT-100M, 768 dimensional for NT-250M, DNABERT-2, and DNABERT-S). To compute mean pooling, we expand the attention mask to match the dimensions of the hidden states, then compute the sum of the token embeddings weighted by the attention mask, and divide by the total number of valid (non-padded) tokens. This ensures only meaningful tokens contribute to the final sequence embedding. Finally, the mean pooled embeddings are passed through each of the classification heads to produce logits corresponding to each hierarchical level. During training, a separate cross-entropy loss is computed for each level, and the overall loss is calculated as the sum of these individual losses. We opted not to use a class-weighted training loss, as we found that incorporating class weights led to more uneven performance across different classes.

### Training and Resources

For all experimental runs (full fine-tuning, random initialization, partial freezing layers, LoRA, and (IA)^3^), the Nucleotide Transformer models (NT-50M, NT-100M, NT-250M) were trained on a single NVIDIA H100 GPU with 80 GB of VRAM. These models were trained with per-device batch sizes of 128, 64, and 64, and corresponding learning rates of 2e-4, 1e-4, and 1e-4, respectively. In contrast, the DNABERT models (DNABERT-2-117M and DNABERT-S) were trained using 4 NVIDIA H100 GPUs with 80 GB VRAM each, with a per-device batch size of 16 and a learning rate of 2e-5 for both models. All models were trained for a maximum of 5 epochs, with early stopping triggered if the validation loss did not improve for 3 consecutive epochs. Training was conducted using FP32 precision, with zero weight decay and no warmup phase. A cosine learning rate scheduler was used to adjust learning rate during training. We explored training PEFT models with warmup, a weight decay of 0.1, additional learning rates of 2e-5 and 1e-4, and as many as 20 epochs to verify we were not underfitting or overfitting. We found no meaningful benefit to any of these changes in training hyperparameters with even an additional 15 epochs resulting in a 1% decrease in loss, and still remained substantially higher than the loss of the fully and partially trained models.

Our motivation for testing randomly initialized weights was primarily to test the claims in Vishniakov, et al. (2024) that pretrained gLMs performed worse than randomly initialized models, a claim that seemed counter to experiments across a number of domains. For our experiments, we reset the weights of all linear and embedding layers using values drawn from a normal distribution with mean zero and a standard deviation specified by the model’s configuration. Layer normalization weights are set to one and biases to zero, while biases in linear layers are also zeroed. This process mirrors the standard initialization of transformer architectures, and discards all pretrained knowledge by allowing the model to be trained from scratch.

